# Functional genetics reveals modulators of anti-microtubule drug sensitivity

**DOI:** 10.1101/2024.03.12.584469

**Authors:** Kuan-Chung Su, Elena Radul, Nolan K Maier, Mary-Jane Tsang, Claire Goul, Brittania Moodie, Heather R. Keys, Iain M Cheeseman

## Abstract

Microtubules play essential roles in diverse cellular processes and are important pharmacological targets for treating human disease. Here, we sought to identify cellular factors that modulate the sensitivity of cells to anti-microtubule drugs. We conducted a genome-wide CRISPR/Cas9-based functional genetics screen in human cells treated with the microtubule-destabilizing drug nocodazole or the microtubule-stabilizing drug taxol. We further conducted a focused secondary screen to test drug sensitivity for ∼1400 gene targets across two distinct human cell lines and to additionally test sensitivity to the Kif11-inhibitor, STLC. These screens defined gene targets whose loss enhances or suppresses sensitivity to anti-microtubule drugs. In addition to gene targets whose loss sensitized cells to multiple compounds, we observed cases of differential sensitivity to specific compounds and differing requirements between cell lines. Our downstream molecular analysis further revealed additional roles for established microtubule-associated proteins and identified new players in microtubule function.

## Introduction

Microtubules are dynamic cytoskeletal polymers that serve as essential structural and force-generating elements in all eukaryotic cells. Microtubule assembly involves heterodimers of α-tubulin and β-tubulin associating in a head-to-tail fashion to form a polarized polymer, termed a protofilament, with each microtubule composed of 11-15 protofilaments (Chaaban et al., 2018). The microtubule minus end is often stabilized or anchored at specific cellular sites, whereas the plus ends can undergo growth and shortening (Chalfie and Thomson, 1982; Cueva et al., 2012; Desai and Mitchison, 1997; Kapitein and Hoogenraad, 2011). Microtubules are highly dynamic, undergoing spontaneous growth, shortening, and regrowth – a behavior termed ‘dynamic instability’ (Mitchison and Kirschner, 1984). These intrinsic features of microtubules are further modulated by a network of microtubule-associated proteins to create differing microtubule behaviors, dynamics, and organization (reviewed in (Akhmanova and Steinmetz, 2015; Bodakuntla et al., 2019)), including creating differences between cell types and over the course of the cell cycle (Chaaban et al., 2018; Howes et al., 2017).

Previously identified microtubule-associated proteins display a range of different properties and activities (Ishikawa, 2017; Karsenti et al., 2006; Lin and Nicastro, 2018; Nedelec et al., 2003). For example, some microtubule-associated proteins bind specifically to the microtubule plus or minus ends to control microtubules dynamics or the facilitate the attachment of microtubules to the cell cortex, kinetochores, intracellular membrane organelles, and other cellular structures. Other microtubule-associated proteins bind to the microtubule lattice to regulate microtubule interactions, organization, dynamics, and stability. Finally, molecular motors, such as dynein and kinesin-related proteins, are responsible for transporting cargos along microtubules or generating forces within microtubule structures such as in the mitotic spindle or the flagellar axoneme.

Microtubules are also important pharmacological targets for treating human disease. During mitosis, even modest alterations to microtubule dynamics can lead to chromosome instability. As a result, mitotic cells are sensitive to anti-microtubule agents, including the microtubule-stabilizing drug taxol and the microtubule-depolymerizing drug nocodazole. Tubulin-targeting drugs, such as taxol and vincristine, are frontline chemotherapeutics against various types of cancer, and the development of resistance to anti-microtubule drugs represents a substantial obstacle in cancer treatment (Cermak et al., 2020; Prassanawar and Panda, 2019; Zhou and Giannakakou, 2005).

Given the central roles for microtubules across diverse cellular processes, defining the complete network of proteins involved in microtubule regulation, dynamics, organization, and function is critical. The use of CRISPR/Cas9-based pooled functional genetic screens has revolutionized the ability to identify the genes, pathways, and mechanisms involved in a biological process by enabling the disruption of thousands of individual genes (Shalem et al., 2014; Wang et al., 2014). In our recent work, we used optical pooled screening to identify essential genes whose depletion results in dramatic alterations to microtubule assembly or organization (Funk et al., 2022). Here, as an complementary strategy to identify genes with roles in the microtubule cytoskeleton that impact cellular fitness, we conducted large-scale CRISPR/Cas9 functional screening in the presence of low doses of either the microtubule-destabilizing drug nocodazole (Vasquez et al., 1997), the microtubule-stabilizing drug taxol (Arnal and Wade, 1995; Schiff and Horwitz, 1980; Yang and Horwitz, 2017), or the KIF11 inhibitor STLC, which disrupts bipolar spindle assembly (Kaan et al., 2009; Wu et al., 2018). These pooled screens allowed us to define regulators of microtubule dynamics that enhance or suppress cell proliferation or survival in the presence of one or more of these drugs. Using this approach, we identified multiple established players in controlling microtubule dynamics and spindle organization as well as additional factors that have not been implicated previously in microtubule function in human cells.

## Results and Discussion

### A genome-wide pooled functional genetic screen reveals modulators of microtubule drug sensitivity

To identify factors that modulate the sensitivity of cells to microtubule-based drugs, we conducted pooled genome-wide CRISPR/Cas9-based screening in cultured human cells (Fig. 1A). For this initial screen, we compared the growth behavior of untreated leukemia-derived K562 cells with cells grown in the presence of either the microtubule-destabilizing drug nocodazole or the microtubule-stabilizing drug taxol (Fig. 1B). To identify both enhancers and suppressors of drug sensitivity, we used non-lethal drug concentrations below the IC50 in which growth was only modestly affected (∼5% reduced proliferation per doubling). We conducted the screen over 14 population doublings, with drug-containing media changed every 2 days. High throughput sequencing of the single guide RNA (sgRNA) representation in the terminal timepoint relative to the population at day 0 allowed us to generate a CRISPR score indicative of the fitness consequence of targeting each gene (Fig. 1A, Table S1) (Shalem et al.; Wang et al., 2014). A negative CRISPR score indicates that the elimination of the corresponding gene results in reduced cell proliferation or survival, with strongly negative scores (< –1) consistent with an essential requirement for the gene under the tested growth conditions. Reciprocally, a positive score indicates that the targeted cells proliferate more quickly or survive more robustly than other cells in the population, which will include suppressors of drug treatment.

**Figure 1:**
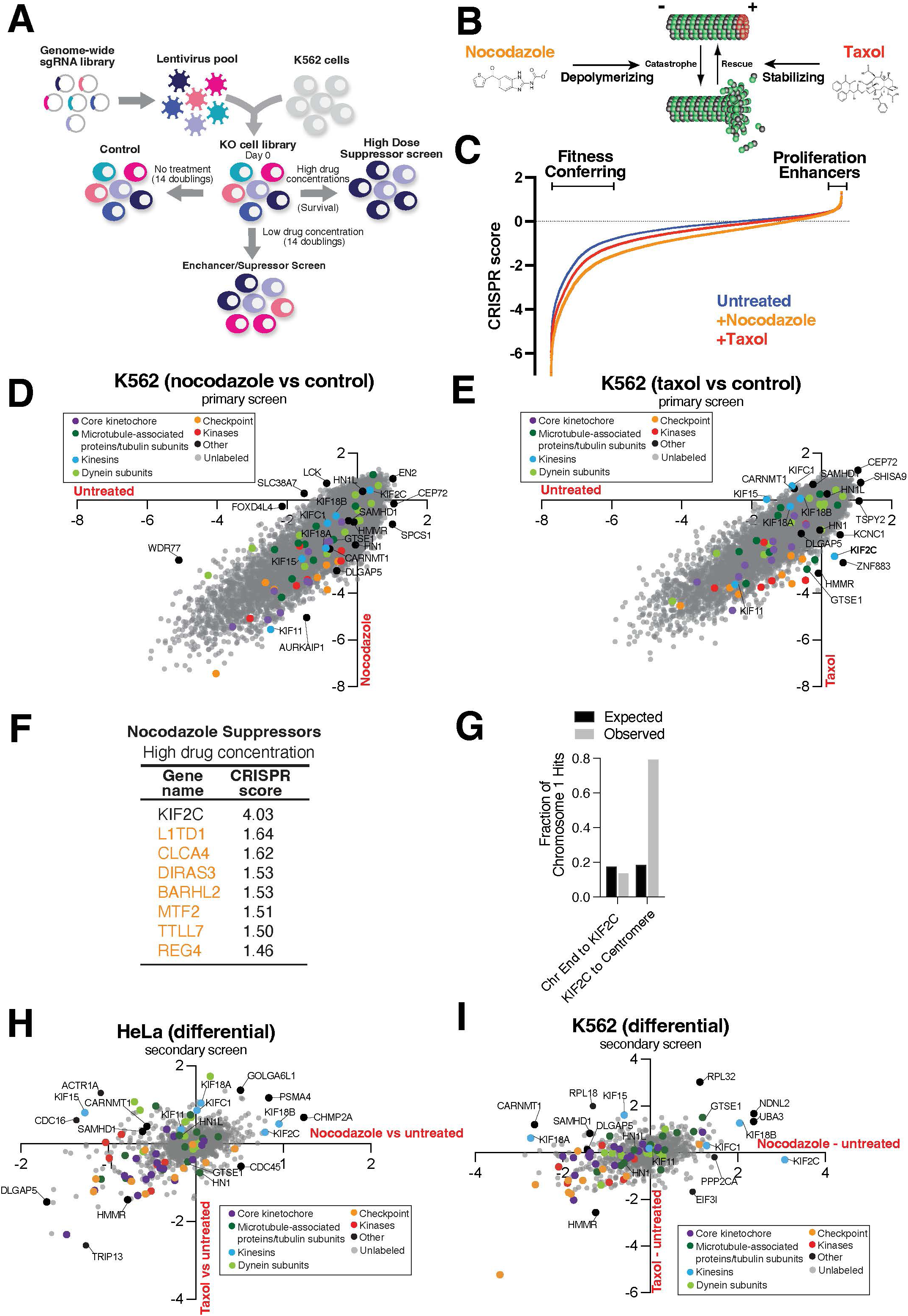
Large-scale functional genetics screens reveal modulators of anti-microtubule drug sensitivity. (A) Schematic showing the workflow for the pooled CRISPR screen. (B) Schematic showing the effects of microtubule drugs (nocodazole and taxol) on microtubule dynamics.(C) Curve illustrating the CRISPR score of fitness conferring and growth enhancer genes upon treatment with either nocodazole or taxol. (D) Scatter plot showing the CRISPR scores in untreated versus nocodazole treated cell pools. (E) Scatter plot showing the CRISPR scores in untreated versus taxol treated cell pools. (F) Table showing the CRISPR scores of selected inducible knockouts (iKO) in high concentration of nocodazole or taxol. (G) Expected versus observed fractions of hits on between the end of chromosome 1 and KIF2C, or between KIF2C and the centromere. Expected fractions were calculated by assuming an even distribution of hits across the chromosome 1 length. (H) Scatter plot illustrating the differential CRISPR scores across all gene targets in the secondary screen. The differential was calculated between nocodazole and untreated and taxol and untreated HeLa cell pools. (I) Scatter plot illustrating the differential CRISPR scores across all gene targets in the secondary screen. The differential was calculated between nocodazole and untreated and taxol and untreated K562 cell pools.

Targeting the vast majority of genes resulted in similar overall fitness effects in both control and drug-treated populations (Fig. 1C-E), with an R^2^ of 0.73 for control vs. nocodazole treated cells and 0.78 for control vs. taxol treated cells. However, a subset of gene targets displayed altered cellular fitness in the presence of microtubule drugs (Fig. 1C-E). This includes diverse factors involved in microtubule-related and cell division-related functions, such as kinetochore proteins, spindle assembly checkpoint components, mitotic kinases, microtubule-associated proteins, and microtubule-based motors. Although many gene targets displayed similar compromised growth in the presence of either taxol or nocodazole, some targets displayed differential behavior between the two drugs (Fig. 1H). Gene targets with substantially altered growth in at least one drug condition were selected for further analysis, as described below.

### Loss of KIF2C/MCAK acts as a dose-sensitive suppressor for growth in nocodazole

In addition to our analysis of drug sensitivity in the presence of modest levels of anti-microtubule drugs, we also conducted a screen for gene targets that were able to confer resistance to increased drug concentrations. To identify such suppressors, we gradually increased the drug concentration during each passage of the cells, ultimately reaching a concentration of 250 nM nocodazole or 15 nM taxol. Based on the change in relative sgRNA abundance in the presence and absence of drug treatment, the most potent suppressor of drug sensitivity was the kinesin-13 family member KIF2C/MCAK, whose loss resulted in a strong suppression of nocodazole sensitivity (Fig. 1F). In contrast, we did not identify clear suppressors of high dose taxol treatment. KIF2C binds to microtubule plus ends to accelerate microtubule depolymerization (Howard and Hyman, 2007; Walczak et al., 2013; Wang et al., 2015). Thus, loss of KIF2C is predicted to result in more stable microtubules and buffer cells from perturbations that induce microtubule disassembly, such as nocodazole. Reciprocally, we observed that KIF2C knockouts displayed enhanced sensitivity at low taxol concentrations (Fig. 1E), consistent with a model in which Kif2C-depleted cells have more stable microtubules and therefore cannot tolerate additional stabilizing treatments.

To identify additional suppressors of nocodazole sensitivity, we surveyed gene targets that were non-essential in untreated cells, but displayed improved fitness in nocodazole. Surprisingly, we observed that the majority of nocodazole suppressors were spatially clustered on chromosome 1. In particular, these suppressors were located between KIF2C and the centromere on the p arm of chromosome 1 with a substantial enrichment of hits compared to a random distribution (Figure S1A). Based on the observed clustering, we hypothesize that targeting these genes with sgRNAs results in Cas9-mediated DNA cleavage, which in the absence of DNA repair would lead to loss of the arm of chromosome 1 containing KIF2C. Although such events likely occur at a low frequency, nocodazole treatment may select for cells with heterozygous loss of the p-arm of chromosome 1 due to the potent suppressive effect of Kif2C-depletion on nocodazole-mediated toxicity. This chromosome arm-loss and suppression behavior has also been observed by others (Ashoti et al., 2022; Cullot et al., 2019) and underscores its relevance for CRISPR screens with potent dose-sensitive suppressors, in which Cas9-mediated chromosome arm cleavage may contribute to false positives. Together, these results demonstrate that the loss of KIF2C acts as a robust suppressor of the microtubule-destabilizing drug nocodazole.

### Secondary screening identifies modulators of drug sensitivity across diverse conditions

To validate the fitness behaviors observed in the primary screen and identify factors that suppress or enhance the anti-proliferative effects of anti-microtubule drug treatment, we next conducted a secondary screen targeting a reduced set of 1411 genes composed of established cell division components, genes that displayed differential growth in nocodazole or taxol in the primary screen (Fig. 1 D-E, S1B), and additional control factors. For this analysis, we conducted CRISPR/Cas9 screens in both the suspension cell line, K562 (Figure 1I), and the adherent human cervical cancer cell line, HeLa (Figure 1H). In each case, we tested the fitness effects of targeting these genes in the presence of low doses of nocodazole, taxol, and the KIF11 inhibitor STLC. These datasets provide rich information across a range of comparisons (Figure S1C-E, S2, Table S2), including identifying enhancers and suppressors of the different drug treatments, gene targets that result in differential effects between drug treatments, and differences between cell lines. Amongst these gene targets and growth properties, we selected genes that displayed unexpected behaviors or had not been previously implicated in microtubule-related functions for in-depth analysis. In particular, we selected two established players in microtubule dynamics, KIF15 and DLGAP5/HURP, and five genes with less established roles (HMMR, SAMDH1, HN1, HN1L and CARNMT1) for further analysis to determine their contributions to microtubule assembly, spindle formation, and chromosome segregation.

### Loss of KIF15 and DLGAP5 lead to the formation of monopolar spindles upon nocodazole treatment

Our pooled Cas9-based screens indicated that loss of the kinesin-12 family member KIF15 (also known as Hklp2; (Drechsler et al., 2014)) and the microtubule-associated protein DLGAP5 (also called HURP) sensitized cells to nocodazole treatment (Fig. 2A). To test the functions of KIF15 and DLGAP5, we conditionally eliminated each gene using a doxycycline-inducible Cas9-based system (McKinley and Cheeseman, 2017; McKinley et al., 2015) to generate inducible knockout (iKO) HeLa cell lines. Based on immunofluorescence analysis, we found that inducible knockout of either KIF15 or DLGAP5 resulted in a modest increase in the frequency of chromosome alignment errors, particularly off-axis chromosomes (Fig. 2B-D).

**Figure 2:**
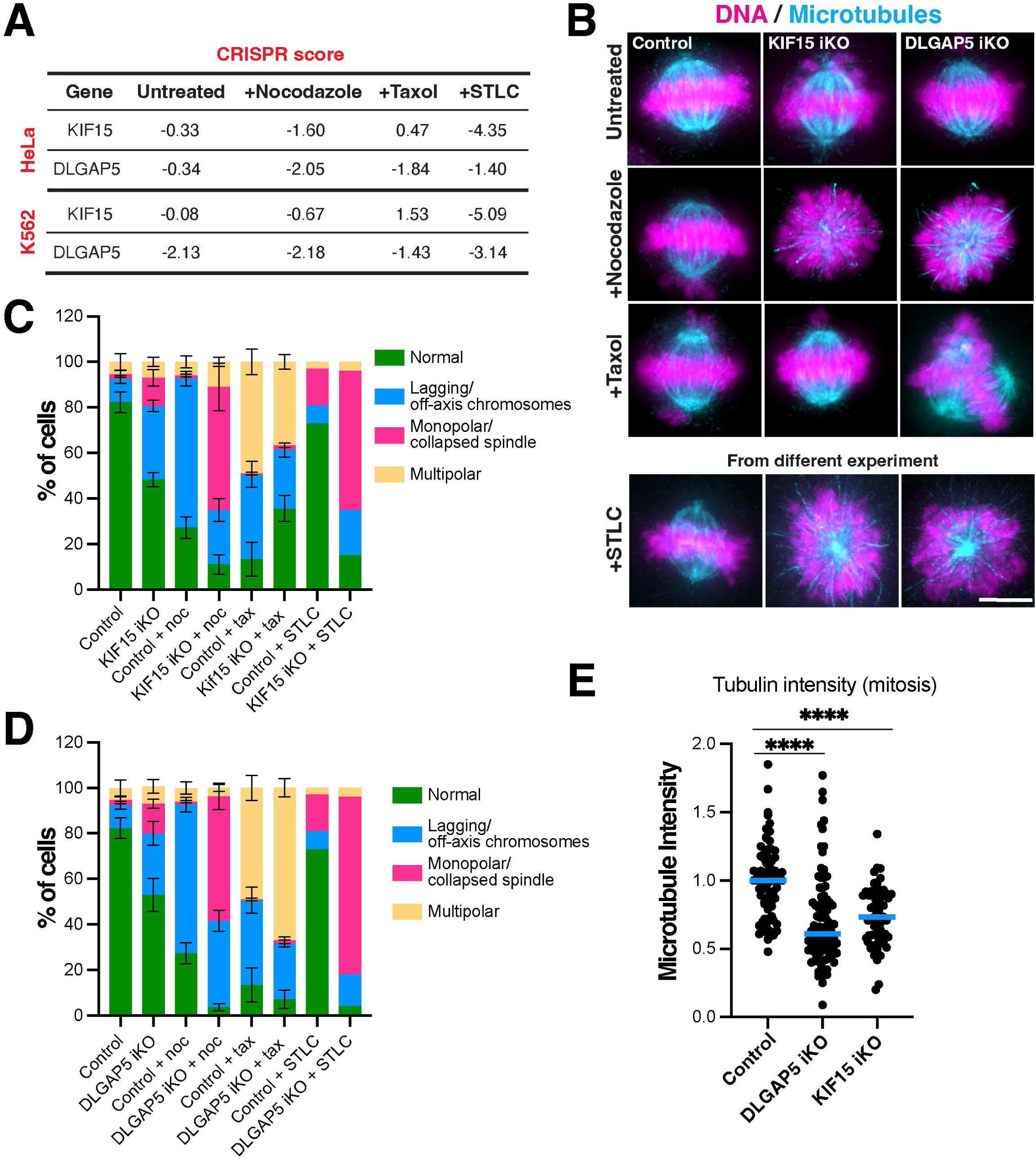
Analysis of KIF15 and DLGAP5 inducible knock-outs. (A) Table showing the secondary screen CRISPR score of KIF15 and DLGAP5 inducible knockouts (iKOs) in HeLa and K562 cells. (B) Representative Z-projected widefield immunofluorescence images of mitotic metaphase cells from KIF15 and DLGAP5 iKO HeLa cell lines treated with either nocodazole, taxol or STLC. Microtubules (DM1α), DNA (DAPI). Scale bar: 9 µm. (C) Percent of mitotic cells with chromosome misalignment defects after iKO of KIF15 for 4 d, treated with either nocodazole, taxol or STLC, quantified from B. n = approximately 300 cells per condition, across three experimental replicates. (D) Percent of mitotic cells with chromosome misalignment defects after iKO of DLGAP5 for 4 d, treated with either nocodazole, taxol or STLC, quantified from B. n = approximately 300 cells per condition, across three experimental replicates. (E) Quantification of total spindle tubulin immunofluorescence in KIF15 iKO and DLGAP5 iKO HeLa cells. n = 71, 60, 96 across three experimental replicates. Statistical tests performed: Welch’s t test (**** = < 0.0001).

Based on the behaviors observed in the functional genetics screens, we next tested the effects of combining KIF15 or DLGAP5 knockouts with anti-microtubule drug perturbations. Interestingly, although low-dose nocodazole only modestly compromised spindle organization on its own, loss of either KIF15 or DLGAP5 led to a ∼50% increase in the frequency of monopolar spindles following nocodazole addition (Figure 2C). In our genome-wide screen, we observed an increased CRISPR score for KIF15 and decreased fitness for DLGAP5 in taxol-treated cells (Fig. 1E, 2A, S2B). Consistent with this differing drug sensitivity, we found that low-dose taxol treatment modestly reduced the frequency of chromosome alignment errors in KIF15 knockout cells, but resulted in an increased proportion of multipolar spindles in DLGAP5 knockout cells (Fig 2B-D). Finally, targeting of either KIF15 or DLGAP5 resulted in a substantially-increased fraction of monopolar spindles upon treatment with STLC compared to control cells. For KIF15, this is in agreement with previous findings in which KIF15 becomes indispensable for spindle formation when KIF11 is perturbed (Drechsler et al., 2014), and its established role in bipolar spindle formation (Tanenbaum et al., 2009). Based on the quantification of total tubulin fluorescence, we found that the microtubule intensity of the mitotic spindle was reduced by 39% in KIF15-depleted cells and 45% in DLGAP5-depleted cells when compared to control cells (Figure 2E, S3A,B). In contrast, we did not observe dramatic defects in microtubule organization or abundance in interphase HeLa cells (not shown).

Taken together, these results validate the KIF15 and DLGAP5 knockout growth behavior observed in the primary and secondary screens (Fig. 1) and show that bipolar spindle formation in these knockouts is highly sensitive to microtubule depolymerization. In addition, the differential growth behavior between nocodazole and taxol treatment observed in Kif15 knockout cells, as well as the reduced microtubule polymer levels, suggest that KIF15 might play roles in modulating microtubule dynamics beyond its established function in spindle pole separation.

### Implication of HMMR and SAMHD1 in microtubule organization

Amongst the additional factors that altered growth in the presence of anti-microtubule drugs, we selected the microtubule-associated hyaluronan-mediated motility receptor (HMMR) protein and the SAM and HD domain-containing protein 1 (SAMHD1) for further analysis. HMMR is a nonmotor adaptor protein that contributes to mitotic spindle assembly by stabilizing kinetochore-fibers (Chen et al., 2018; Manning and Compton, 2008). SAMHD1 is a dNTP triphosphohydrolase that plays a crucial role during viral infection (Beloglazova et al., 2013; Goldstone et al., 2011; White et al., 2013), but has not been implicated previously in microtubule dynamics. In the pooled screens, loss of HMMR sensitized cells to both nocodazole and taxol treatment, whereas loss of SAMHD1 sensitized cells to nocodazole and STLC treatment (Fig. 3A, C).

**Figure 3:**
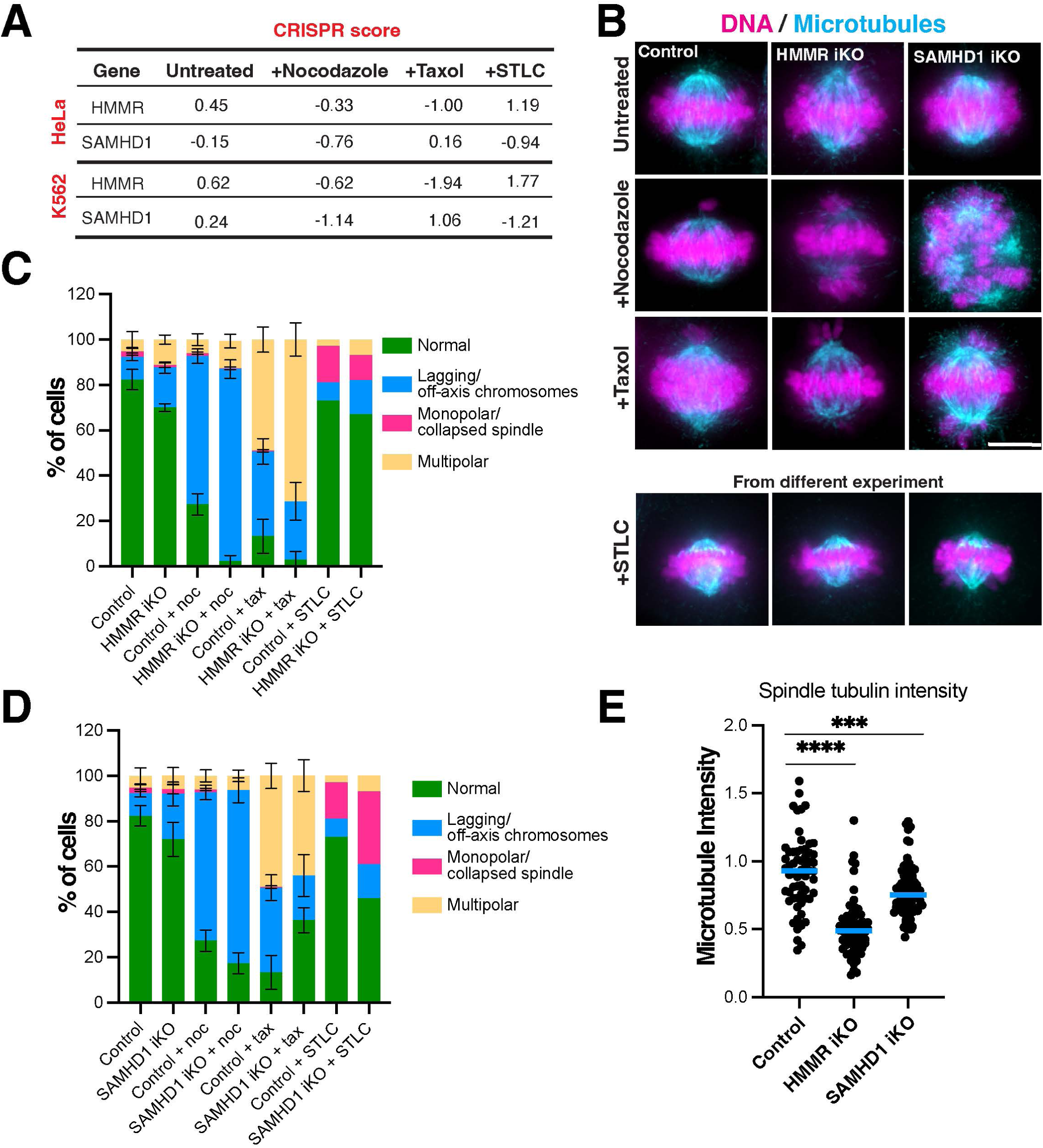
Analysis of: HMMR and SAMHD1 inducible knock-outs. (A) Table showing the secondary screen CRISPR score of HMMR and SAMHD1 inducible knockouts (iKOs) in HeLa and K562 cells. (B) Representative Z-projected widefield immunofluorescence images of mitotic metaphase cells from HMMR and SAMHD1 iKO HeLa cell lines treated with either nocodazole, taxol or STLC. Microtubules (DM1α), DNA (DAPI). Scale bar: 9 µm. (C) Percent of mitotic cells with chromosome misalignment defects after iKO of HMMR for 4 d, treated with either nocodazole, taxol or STLC, quantified from B. n = approximately 300 cells per condition, across three experimental replicates. (D) Percent of mitotic cells with chromosome misalignment defects after iKO of SAMHD1 for 4 d, treated with either nocodazole, taxol or STLC, quantified from B. n = approximately 300 cells per condition, across three experimental replicates. (E) Quantification of total spindle tubulin immunofluorescence in the HMMR and SAMHD1 iKO HeLa cells. n = 61, 69, 62 across three experimental replicates. Statistical tests performed: Welch’s t test (**** = < 0.0001).

In inducible knockout cell lines, we found that HMMR knockout led to a ∼20% increase in the frequency of off-axis chromosomes, consistent with prior reports (Chen et al., 2018). Treatment of HMMR iKO cells with low dose nocodazole caused a further ∼20% increase in cells with off-axis chromosomes and other spindle defects. However, we observed the most dramatic effect upon treatment with low dose taxol, which led to ∼20% increase in the proportion of cells with multipolar spindles, many with 4-5 spindle poles (Figure 3C). In SAMHD1 knockouts, we observed a reduction in lagging chromosomes upon taxol treatment, but no significant changes in the frequency of chromosome segregation errors in nocodazole-treated cells (Figure 3B, C). Finally, when quantifying total tubulin intensity, we observed that the microtubule mass on the spindle was reduced by 41% in HMMR-depleted cells (Figure 3E), and we observed a small, but significant tubulin intensity difference in SAMHD1-depleted cells when compared to controls (Fig. 3E; Fig. S3D,E). These results highlight contributions of HMMR and SAMHD1 to mitotic spindle function, activities that are particularly revealed in the presence of modest perturbations to microtubule dynamics. We also note that SAMHD1 does not show apparent localization to microtubule structures (Fig. S3C), suggesting that it acts indirectly to influence microtubule behavior.

### HN1L-HN1 double knockout increases nocodazole sensitivity and taxol resistance

In the primary screen, targeting of the hematological and neurological expressed 1 (HN1; also known as JPT1) led to substantially reduced growth in the presence of both taxol and nocodazole (Fig. 4A). However, we did not observe similarly strong phenotypes for HN1-targeted cells in the secondary screen (Fig. 4A). We reasoned that this experimental variability could reflect partial redundancy between HN1 and HN1L (also known as JPT2). We found that GFP fusions with either HN1 or HN1L localized to both interphase microtubules and mitotic spindle microtubules (Fig. 4B). This is consistent with the reported microtubule localization for the *Drosophila* HN1/HNL1 homolog Jupiter (Karpova et al., 2006), but this microtubule association had not been tested previously in human cells.

**Figure 4:**
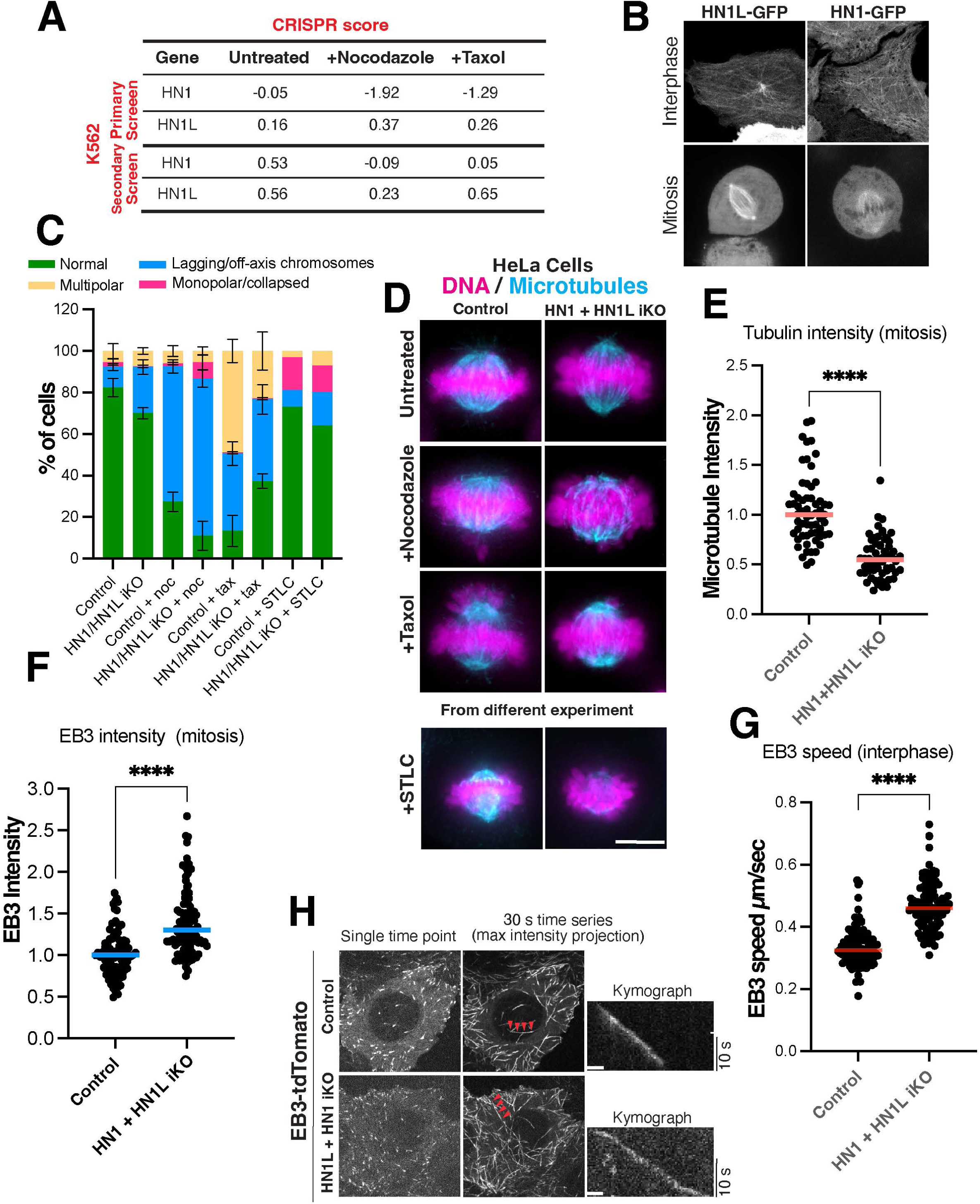
Analysis of HN1 and HN1L double knock-out cells. (A) Table showing the primary and secondary screen CRISPR score of HN1 and HN1L inducible knockouts (iKOs) in K562 cells. (B) Representative confocal immunofluorescence images of mitotic metaphase and interphase HeLa cells showing the localization of GFP-tagged HN1 and HN1L proteins. (C) Percent of mitotic cells with chromosome misalignment defects after iKO of HN1 and HN1L for 4 d, treated with either nocodazole, taxol or STLC, quantified from D. n = approximately 300 cells per condition, across three experimental replicates. (D) Representative Z-projected widefield immunofluorescence images of mitotic metaphase cells from HN1/HN1L double iKO HeLa cell line treated with either nocodazole, taxol or STLC. Microtubules (DM1α), DNA (DAPI). Scale bar: 9 µm. (E) Quantification of total spindle tubulin immunofluorescence in the HN1 and HN1L iKO HeLa cells. n = 60, 62 across three experimental replicates. (F) Quantification of total spindle EB1 immunofluorescence in the HN1 and HN1L iKO HeLa cells. n = 94, 105 across three experimental replicates. (H) Live confocal immunofluorescence images of td-Tomato EB3 tagged HN1 and HN1L iKO HeLa cells. (G) EB3 speed quantification in HN1 and HN1L iKO HeLa cells. n = 108, 103 kymographs, n = 40, 42 cells across three experimental replicates. Statistical tests performed: Welch’s t test (**** = < 0.0001).

Knockout of either HN1 or HN1L alone did not result in increased chromosome segregation defects or altered microtubule intensity (Fig. S4A). Thus, to test for redundancy between HN1 and HN1L, we simultaneously eliminated both proteins. We found that HN1+HN1L double knockouts displayed a mild increase in chromosome segregation defects compared to control cells (Fig. 4C, D). However, upon treatment with a low dose of nocodazole, the fraction of cells with chromosome segregation defects increased by 20% (Fig. 4C). Reciprocally, HN1+HN1L double knockout cells treated with a low dose of taxol displayed a significant decrease in the fraction of multipolar cells. To test the basis for these phenotypes, we analyzed the consequences to microtubule dynamics. We observed that the microtubule mass on the spindle was reduced by 52% in HN1+HN1L-depleted cells, suggesting that microtubules are destabilized in these knockout cells. In addition, the amount of plus-end tracking protein EB3, a proxy for the relative level of microtubule dynamics on the spindle (Akhmanova and Steinmetz, 2008; Nehlig et al., 2017; Telley et al., 2011) was increased by 76% (Fig. 4E, F). These behaviors suggest that the combined loss of HN1 and HN1L leads to increased spindle microtubule dynamics. In interphase HeLa cells, the HN1+HN1L double knockout led to a 39% increase in the microtubule polymerization rate based on the imaging of EB3 (Fig. 4H, G S4B). Nevertheless, we also observed a modest increase in mitotic tubulin signal (Fig. S4C,D). Together, our results suggest that the simultaneous loss of both HN1 and HN1L leads to more dynamic microtubules in both interphase and mitotic cells.

### CARNMT1 promotes microtubule stability

As both nocodazole and taxol disrupt chromosome segregation, many gene knockouts displayed similar sensitivity to treatment with both of these compounds (Fig. 1H,I). However, as these drugs have opposing effects on microtubule stability, factors that modulate microtubule dynamics are predicted to have different growth effects in the presence of these drugs. For example, we found that the microtubule-destabilizing motor KIF2C/MCAK suppressed nocodazole treatment, but modestly enhanced taxol treatment in K562 cells (Fig. 1I). We therefore sought to analyze other gene targets that displayed differential growth behaviors. The loss of carnosine N-methyltransferase 1 (CARNMT1 or C9orf41) led to opposing growth behavior in nocodazole and taxol, with increased fitness in taxol-treated cells and decreased fitness in nocodazole-treated cells (Fig. 5A). From these data, we predicted that CARNMT1 acts to stabilize microtubules.

**Figure 5:**
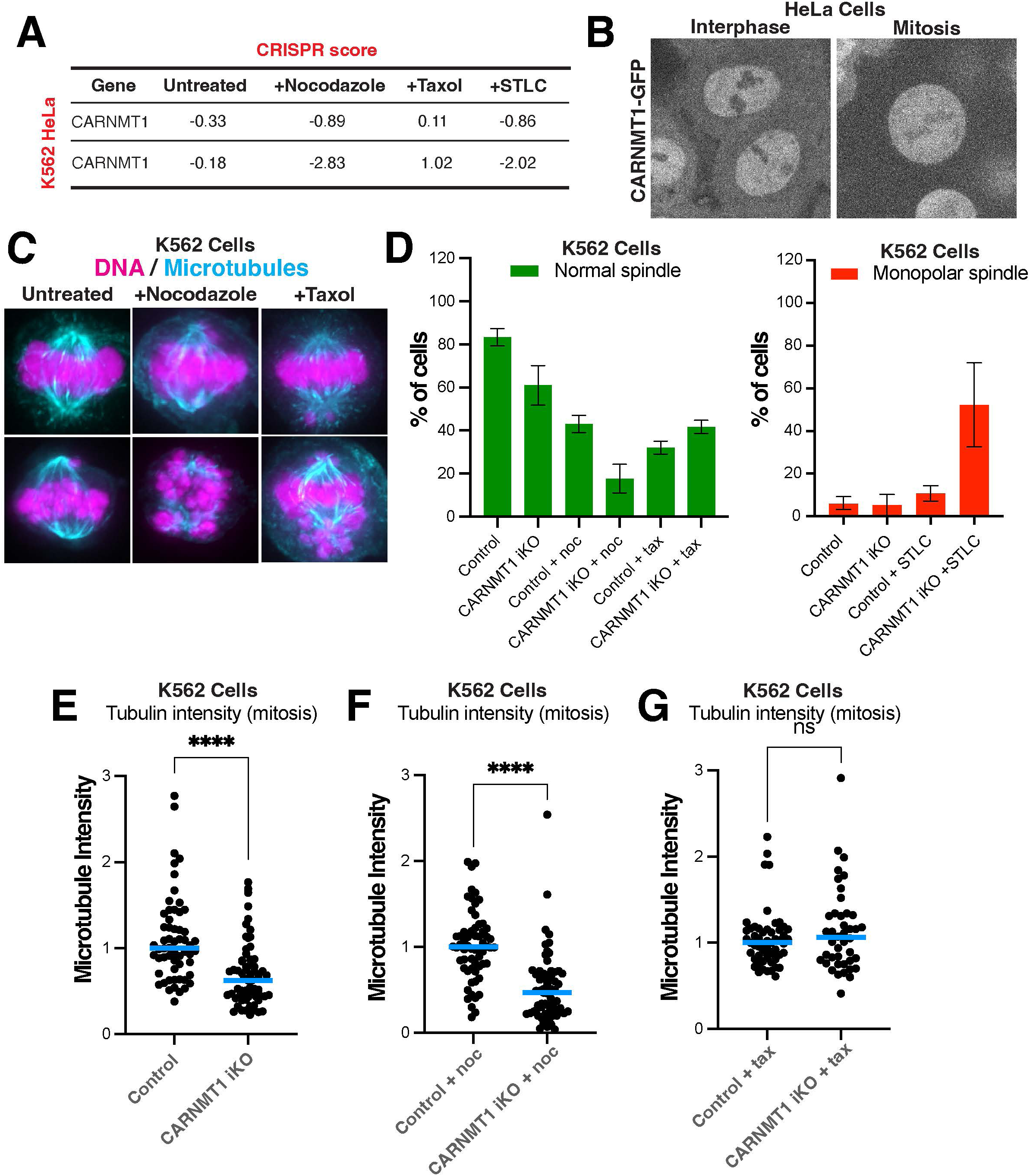
Analysis of CARNMT1 knock-out cells. (A) Table showing the secondary screen CRISPR score of CARNMT1 iKO in HeLa and K562 cells. (B) Representative confocal immunofluorescence images of mitotic metaphase and interphase HeLa cells showing the localization of GFP-tagged CARNMT1. (C) Representative Z-projected widefield immunofluorescence images of mitotic metaphase cells from CARNMT1 iKO K562 cell line treated with either nocodazole, taxol or STLC. Microtubules (DM1α), DNA (DAPI). Scale bar: 9 µm. (D) Percent of mitotic cells with chromosome misalignment defects after iKO of CARNMT1 for 4 d, treated with either nocodazole, taxol or STLC, quantified from C. n = approximately 300 cells per condition, across three experimental replicates. (E) Quantification of total spindle tubulin immunofluorescence in the CARNMT1 iKO K562 cells. n = 58, 64 across three experimental replicates. (F) Quantification of total spindle tubulin immunofluorescence in the CARNMT1 iKO K562 cells treated with nocodazole. n = 62, 63 across three experimental replicates. (G) Quantification of total spindle tubulin immunofluorescence in the CARNMT1 iKO K562 cells treated with taxol. n = 62, 61 across three experimental replicates. Statistical tests performed: Welch’s t test (**** = < 0.0001).

Prior work found that CARNMT1 is responsible for synthesizing anserine by methylation of carnosine (Cao et al., 2018). Both carnosine and anserine are abundant dipeptides in vertebrate skeletal muscles and are suggested to serve as proton buffers and radical scavengers (Drozak et al., 2015). CARNMT1 localizes to the nucleus in interphase HeLa cells (Fig. 6B) and is diffuse throughout the cell in mitotic cells (not shown), suggesting that it does not associate with microtubules directly. In addition, based on immunoprecipitation experiments, we were unable to identify robust interaction partners for CARNMT1 (data not shown). Thus, if CARNMT1 acts to alter microtubule dynamics, it would function indirectly instead of directly associating with microtubule polymers.

In HeLa cells, CARNMT1 knockouts did not display dramatic changes in the frequency of chromosome segregation errors or changes in the microtubule polymerization rate in interphase cells (Fig. S4E). In our secondary screen, CARNMT1 depletion in K562 cells led to a stronger sensitization to nocodazole than CARNMT1 depletion in HeLa cells (Fig. 5A). Therefore, we generated CARNMT1 inducible knockouts in K562 cells. In K562 cells, we observed that the loss of CARNMT1 led to a modest increase in chromosome segregation defects in comparison to controls, including the presence of chromosome bridges and lagging chromosomes and increased sensitivity to STLC (Fig. 5C,D). CARNMT1 loss also resulted in a modest decrease in the microtubule mass in the spindle (Fig. 5E). Importantly, these segregation defects were enhanced in the presence of low doses of nocodazole with the majority of cells displaying highly scattered chromosomes and a 50% decrease in the total spindle microtubule mass (Fig. 5C-F). Reciprocally, CARNMT1 overexpression led to a slight increase in microtubule mass in the spindle (data not shown). Overall, these results suggest that CARNMT1 acts indirectly to influence microtubule dynamics by acting as a net microtubule-stabilizing factor.

### Identification of Factors that Modulate Microtubule Behavior

Prior studies on microtubule-associated proteins have mostly focused on loss-of-function phenotypes that result in dramatic changes in microtubule assembly and organization (Lindwall and Cole, 1984; Maiato et al., 2003; Mandelkow et al., 1995; Ramkumar et al., 2018). Such approaches are valuable, but likely missed molecular players with non-essential roles in spindle formation that do not result in dramatic defects on their own. Such proteins would still play important roles in microtubule function under specific physiological circumstances, such as in the presence of acute cellular stress, in specific cell types, or following treatment with anti-microtubule compounds such as those that are routinely used in cancer treatment. Our functional genetic studies provide a resource of gene targets whose loss has the potential to modulate the sensitivity to altered microtubule dynamics. Using these approaches, we identified contributions for several established and novel players to spindle function in human cells. In addition, by analyzing cellular phenotypes in a sensitized background using low doses of anti-mitotic compounds, this allowed us to reveal contributions to spindle formation and chromosome segregation that would be missed in an unperturbed background. Importantly, the requirements for these factors and others vary between cell lines and across different anti-microtubule drugs, highlighting the diversity of cellular pathways that contribute to robust spindle function and cell division. Together, our large-scale analysis provides an important resource for considering mitotic spindle function and the factors that may modulate chemotherapeutic drug treatment.

## Acknowledgements

We thank the members of the Cheeseman lab for their support and input. This work was supported by grants from The Harold G & Leila Y. Mathers Charitable Foundation, NSF (2029868) and NIH/NIGMS (R35GM126930) to IMC, a Hope Funds for Cancer Research fellowship (HFCR-18-03-02) to MJT, and the Henry and Frances Keany Rickard Fund Fellowship from the MIT Office of Graduate Education to NKM.

## Author Contributions

Conceptualization KCS, MJT, HRK, IMC; Methodology KCS, ER, NKM, MJT, CG, BM, HRK, IMC, Validation KCS, ER, NKM, MJT, CG, BM, HRK, IMC, Investigation KCS, ER, NKM, MJT, CG, BM, HRK; Writing – Original Draft Preparation KCS, ER, IMC; Writing – Review & Editing KCS, ER, NKM, MJT, CG, HRK, IMC; Visualization ER CG; Supervision: KCS IMC; Funding Acquisition: IMC

## Methods

### Cell culture

HeLa cells (transformed human female cervical epithelium) were cultured in Dulbecco’s modified Eagle medium supplemented with 10% fetal bovine serum, 100 U/mL penicillin and streptomycin, and 2 mM L-glutamine at 37 °C with 5% CO_2_. Doxycycline inducible cell lines were cultured in medium containing fetal bovine serum (FBS) certified as tetracycline free and were induced by addition of doxycycline hyclate to 1 µg/mL for 4 days. Other drugs used on HeLa cells were nocodazole (40 nM; Sigma-Aldrich), paclitaxel (taxol, 5 nM; Invitrogen) and STLC (300 nM; Sigma-Aldrich). The nocodazole and taxol drug treatment was performed for 3 hours whereas the STLC treatment was for 16-17 hours. Hela cells were regularly monitored for mycoplasma contamination using commercial detection kits.

K562 cells were cultured in RPMI-1640 medium supplemented with 10% fetal bovine serum, 100 U/mL penicillin and streptomycin at 37 °C with 5% CO_2_. Doxycycline inducible cell lines were cultured in medium containing FBS certified as tetracycline free and were induced by addition of doxycycline hyclate to 1 µg/mL for 4 days. Other drugs used on these cells were nocodazole (25 nM) and taxol (5 nM). The nocodazole and taxol drug treatment was done for 3 hours while the STLC (350 nM) treatment for 16-17 hours. K562 cells were regularly monitored for mycoplasma contamination using commercial detection kits.

### Pooled CRISPR screens

The specific details for the CRISPR screen are included below. For an extended protocol, see also (Adelmann et al., 2019)

### Generation of lentiviral sgRNA libraries

A validation library comprising 14,989 unique sgRNA sequences targeting 1,406 genes was constructed (Table S1). Genes were chosen based on their differential growth in the initial genome-wide CRISPR screen (either synthetic lethal interactions or increased fitness) and also included established cell division components, and all sgRNAs from an existing genome-wide library (Addgene # 1000000100) targeting each gene in the subset were included in the validation library (10 sgRNAs for most genes). 975 non-targeting control sgRNAs were included, as well as four targeting control sgRNAs resulting in defined numbers of double-strand breaks (CTRL1-AAVS1, CTRL1-HS1, CTRL1-HS15, and CTRL-HS4 (van den Berg et al., 2018)). An upstream adapter was prepended (5’ – TATCTTGTGGAAAGGACGAAACACC – 3’) and a downstream adapter was appended (5’ – GTTTAAGAGCTATGCTGGAAACAGCATAGC – 3’). The library was synthesized as an oligo pool (Agilent). 100 fmol of library per 50 µL PCR reaction was amplified using Q5 HotStart DNA Polymerase (New England Biolabs M0493S) in 8 reactions with a 50 °C – 62 °C gradient annealing step using the following program:

1 cycle 98 °C 2 min

16 cycles 98 °C 10 sec

50 °C – 62 °C 15 sec

72 °C 15 sec

1 cycle 72 °C 2 min

1 cycle 10 °C holdAn aliquot of each reaction was visualized on a 2% agarose gel, and all reactions with the appropriate molecular weight product were combined and cleaned using the DNA Clean and Concentrator 5 kit (Zymo Research D4013). 10 µg LentiCRISPRv2-opti (Addgene #163126) was digested and dephosphorylated for 3 hours in a 60 µL reaction at 37 °C with FastDigest Esp3I and FastAP (ThermoFisher FD0454 and EF0654, respectively). Digested DNA was cleaned using the DNA Clean and Concentrator 5 kit. Insert and digested vector (5 ng: 100 ng per 20 µL reaction) were assembled in 3 separate reactions for 15 minutes at 50 °C or 55 °C using the NEBuilder HiFi DNA Assembly Master Mix (New England Biolabs E2621S) alongside a control reaction omitting the insert. Background assembly was measured by transformation of NEB 5-alpha competent cells (New England Biolabs C2987I) before cleanup of the assembly reactions using AmpureXP magnetic beads (Beckman Coulter A63880). Cleaned assembly reactions were electroporated into Endura Electrocompetent cells (Biosearch Technologies 60242-1) according to manufacturer’s instructions and plated on LB-Lennox 250 mm x 250 mm square bioassay dishes supplemented with 75 µg/mL carbenicillin. Serial dilutions of each electroporation were plated to estimate library coverage. Cells were incubated overnight at 30 °C, collected, and plasmid DNA was prepared using the ZymoPURE II Plasmid DNA Maxiprep kit (Zymo Research D4202). Three electroporations were mixed proportionally to their electroporation efficiency for a combined total library coverage of ∼27-fold. High-throughput sequencing libraries were prepared as in the sequencing library preparation section below using the secondary forward amplification and Read 1 sequencing primers and common reverse amplification and Index sequencing primers. Cycle number was lowered to 14 cycles, and 10 ng template was used per 50 µL reaction.

### Lentivirus Production

For large-scale lentiviral preparation, HEK-293T cells were seeded at a density of 750,000 cells/mL per 175 cm^2^ flask in 20 mL DMEM (Thermo Fisher Scientific) supplemented with 10% FBS (GeminiBio #100-106) and 100U/mL penicillin streptomycin. After 24 hours, media was changed to viral production medium: IMDM (Thermo Fisher Scientific #1244053) supplemented with 20% inactivated fetal serum (GeminiBio #100-106). At 32 hours post-seeding, cells were transfected with a mix containing 76.8 µL Xtremegene-9 transfection reagent (Sigma Aldrich #06365779001), 3.62 µg pCMV-VSV-G (Plasmid #8454, Addgene), 8.28 µg psPAX2 (Plasmid #12260, Addgene), and 20 µg sgRNA plasmid and Opti-MEM (Thermo Fisher Scientific #11058021) to a final volume of 1 mL. Media was changed 16 hours later to fresh viral production medium. At 48 hours after transfection, virus was collected and filtered through a 0.45 µm filter, aliquoted, and stored at –80 °C until use.

### Genome-Wide CRISPR screen

Lentivirus containing the genome-wide lentiviral sgRNA library (Addgene # 1000000100) was used to transduce 500 million K562 cells as previously described (Adelmann et al., 2019) in order to maintain 1000-fold coverage of the sgRNA library. Cells were passaged into fresh RPMI-1640 complete medium supplemented with puromycin (3 µg/ml) at 2 days post-transduction and selected for 3 days. Pellets of 100 million cells were frozen at the endpoint of puromycin selection to assess initial sgRNA library representation. Cells were allowed to recover without puromycin for 2 days before being passaged every 2 days for 14 population doublings under the indicated condition (untreated, low nocodazole (25 – 30 nM), low taxol (2.5 – 3 nM), high nocodazole (increasing concentrations from 37.5 nM to 250 nM), and high taxol (increasing concentrations from 3.75 nM to 15 nM). Pellets of 100 million cells (untreated, low nocodazole, and low taxol) or 5 million cells (high nocodazole and high taxol) were frozen at the endpoint of the screen to assess final sgRNA library representation under each condition.

### Secondary CRISPR screens

For secondary screening of K562 cells, lentivirus containing the targeted lentiviral sgRNA library (Table S2) was used to transduce 35 million K562 cells. Cells were passaged into fresh RPMI-1640 complete medium supplemented with puromycin (3 µg/ml) at 2 days post-transduction and selected for 6 days. Pellets of 5 million cells were frozen at the endpoint of puromycin selection to assess initial sgRNA library representation. Cells were passaged every 2 days for 14 population doublings under the indicated condition (untreated, low nocodazole (15 – 30 nM), low taxol (1.5 – 3 nM), or STLC (500 – 750 nM). Pellets of 5 million cells were frozen at the endpoint of the screen to assess final sgRNA library representation under each condition.

For secondary screening of HeLa cells, lentivirus containing the targeted lentiviral sgRNA library was used to transduce 39 million HeLa cells. Cells were passaged into fresh DMEM complete medium supplemented with puromycin (0.4 µg/ml) at 2 days post-transduction and selected for 4 days. Pellets of 5 million cells were frozen at the endpoint of puromycin selection to assess initial sgRNA library representation. Cells were passaged every 2 days for 14 population doublings under the indicated condition (untreated, low nocodazole (15 – 25 nM), low taxol (1.0 – 1.25 nM), or STLC (125 – 750 nM). Pellets of 5 million cells were frozen at the endpoint of the screen to assess final sgRNA library representation under each condition.

### Sequencing library preparation

From pellets of 100 million cells genomic (g)DNA was extracted using the QIAamp DNA Blood Maxiprep Kit (Qiagen) according to manufacturer’s instructions with the following modifications: 500 µL of a 10 mg/mL solution of ProteinaseK (MilliporeSigma #311587001) in water was used in place of QIAGEN Protease; incubation with ProteinaseK and Buffer AL was performed overnight; centrifugation steps after Buffer AW1 and AW2 were performed for 2 min and 5 min, respectively; gDNA was eluted for 5 min using 1 mL of water preheated to 70 °C, followed by centrifugation for 5 min. Pellets of 5 million cells were extracted using the Blood genomicPrep mini spin kit (Cytiva) according to manufacturer’s instructions, except that cells were lysed at 56 °C overnight, and gDNA was eluted twice consecutively with 30 µL of water preheated to 70 °C. gDNA concentration was determined using the Qubit dsDNA HS Assay kit (Thermo Fisher Scientific #Q32851).

All PCRs were performed in 50 µL reactions using ExTaq Polymerase (Takara Bio #RR001B) with the following program:

1 cycle 95 °C 5 min

28 cycles 95 °C 10 sec

60 °C 15 sec

72 °C 45 sec

1 cycle 72 °C 5 min

1 cycle 4 °C hold

Using the following primers:

Forward (genome-wide): 5’-AATGATACGGCGACCACCGAGATCTACACGAATACTGCCATTTGTCTCAAGATCTA –3’ Forward (secondary): 5’ – AATGATACGGCGACCACCGAGATCTACACCCCACTGACGGGCACCGGA – 3’ Reverse: 5’-CAAGCAGAAGACGGCATACGAGATCnnnnnnTTTCTTGGGTAGTTTGCAGTTTT-3’ Where “nnnnnn” denotes the barcode used for multiplexing.

For all samples, 1, 3, or 6 µg of gDNA was initially amplified for 28 cycles in 50 µL test PCR reactions. For the initial and depletion screen samples, an additional 300 µg of gDNA was used in 50 reactions per sample. For enrichment screens, 27 µg gDNA was subsequently amplified using 6 µg input per reaction. Reactions were pooled and 200 µL of each reaction was purified using AMPure XP magnetic beads (Beckman Coulter # A63880), eluted with 20 µL water, and quantified using the Qubit dsDNA HS Assay kit prior to sequencing for 50 cycles on an Illumina Hiseq 2500 using the following primers: Read 1 sequencing primer (genome-wide): 5’-CGGTGCCACTTTTTCAAGTTGATAACGGACTAGCCTTATTTTAACTTGCTATTTCTA GCTCTAAAAC –3’ Read 1 sequencing primer (secondary): 5’ – GTTGATAACGGACTAGCCTTATTTAAACTTGCTATGCTGTTTCCAGCATAGCTCTTA AAC – 3’ Index sequencing primer: 5’-TTTCAAGTTACGGTAAGCATATGATAGTCCATTTTAAAACATAATTTTAAAACTGCAA ACTACCCAAGAAA-3’

### Cell line generation

The cell lines used in this study are described in table 1. Retrovirus was generated by transfecting 2.5 µg of VSVG packaging plasmid and 5 µg pBABE-based vectors (described in Table 2) into 4 million HEK293-GP cells in 300 µl Buffer EC with 16 µl Enhancer and 60 µl Effectene Transfection Reagent (Qiagen 301425) (Morgenstern and Land, 1990). Supernatant-containing retrovirus was sterile filtered, supplemented with 20 µg/mL polybrene (Millipore) and used to transduce HeLa and K562 containing inducible Cas9. Transduced HeLa and K562 cells were selected with 250 µg/ml hygromycin (Invitrogen) to generate polyclonal cell lines. HeLa and K562 expressing td-Tomato EB3 (pER2) monoclonal cell lines were generated by FACS from the polyclonal cell lines.

**Table 1.**
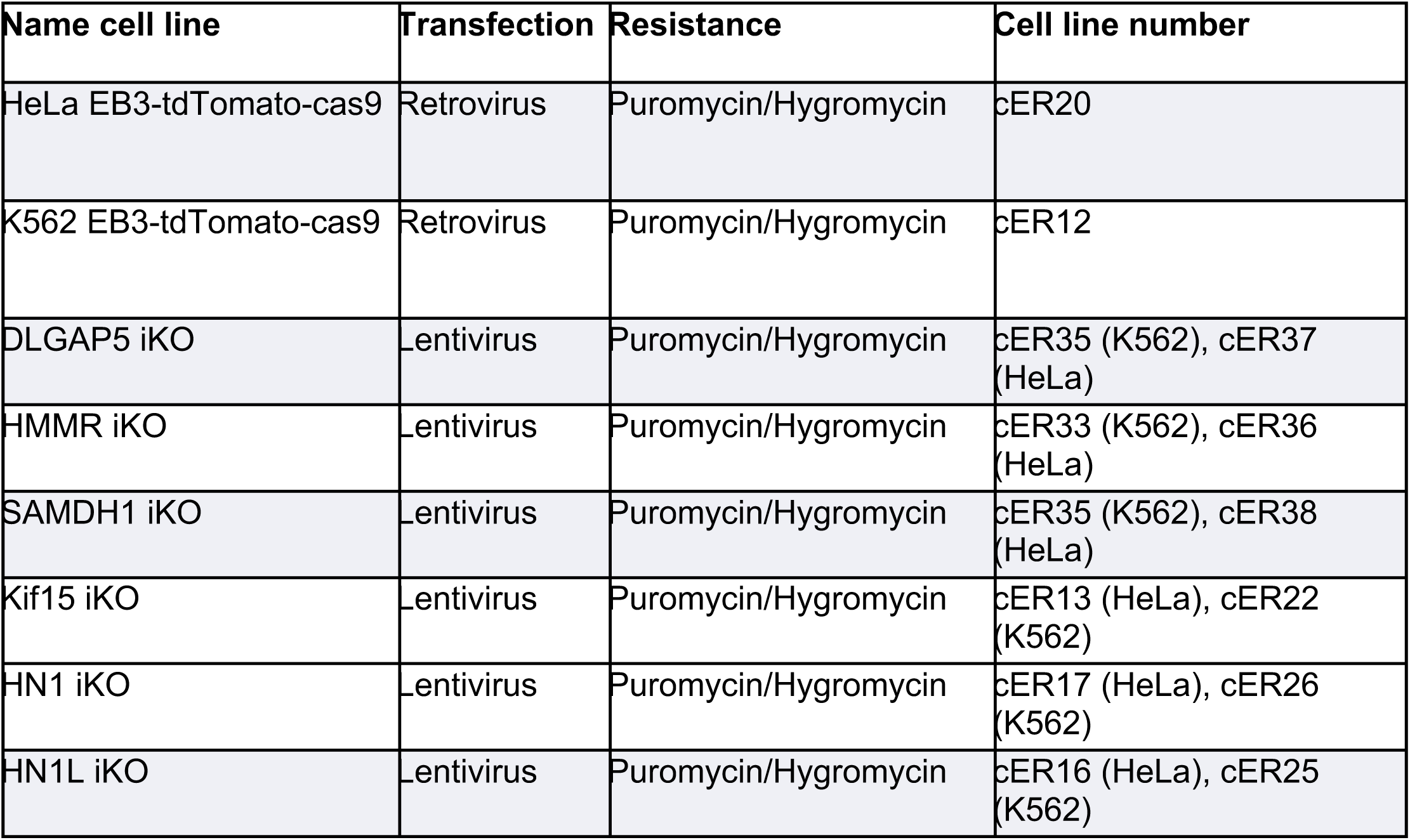
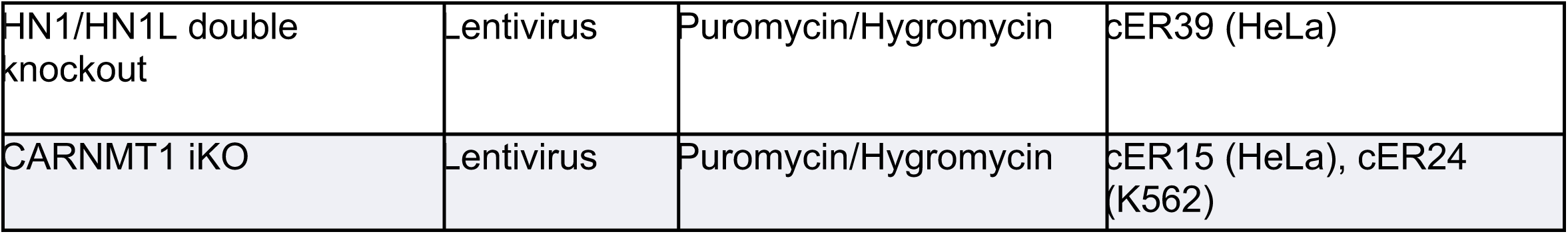
List of cell lines.

| Name cell line | Transfection | Resistance | Cell line number |
| --- | --- | --- | --- |
| HeLa EB3-tdTomato-cas9 | Retrovirus | Puromycin/Hygromycin | cER20 |
| K562 EB3-tdTomato-cas9 | Retrovirus | Puromycin/Hygromycin | cER12 |
| DLGAP5 iKO | Lentivirus | Puromycin/Hygromycin | cER35 (K562), cER37 (HeLa) |
| HMMR iKO | Lentivirus | Puromycin/Hygromycin | cER33 (K562), cER36 (HeLa) |
| SAMDH1 iKO | Lentivirus | Puromycin/Hygromycin | cER35 (K562), cER38 (HeLa) |
| Kif15 iKO | Lentivirus | Puromycin/Hygromycin | cER13 (HeLa), cER22 (K562) |
| HN1 iKO | Lentivirus | Puromycin/Hygromycin | cER17 (HeLa), cER26 (K562) |
| HN1L iKO | Lentivirus | Puromycin/Hygromycin | cER16 (HeLa), cER25 (K562) |
| HN1/HN1L double knockout | Lentivirus | Puromycin/Hygromycin | cER39 (HeLa) |
| CARNMT1 iKO | Lentivirus | Puromycin/Hygromycin | cER15 (HeLa), cER24 (K562) |

**Table 2.**
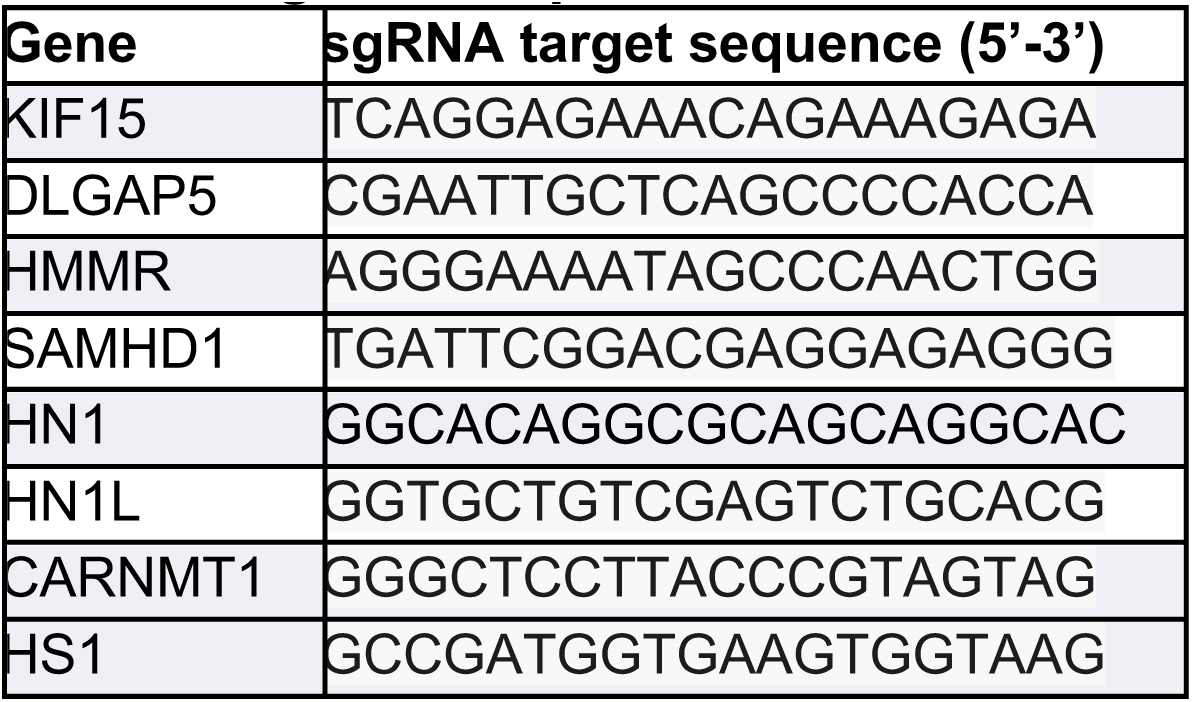
sgRNA sequences.

Lentiviral production and transduction were performed as described previously (McKinley, 2018; McKinley and Cheeseman, 2017; McKinley et al., 2015). Briefly, HEK293FT cells were seeded into 15-cm dishes at a density of 100,000 cells/cm^2^. After one day, cells were transfected with pMD2.G (Addgene #12259), psPAX2 (Addgene #12260), and a lentiviral transfer plasmid (2:3:4 ratio by mass) using Lipofectamine 3000 (Thermo Fisher Scientific L3000015). Viral supernatant was harvested 48h after transfection and filtered through 0.45 µm cellulose acetate filters (Corning 431220). Transduced HeLa and K562 cells were selected with puromycin (Invitrogen) or blasticidin (Invitrogen) as indicated in Table 1 to generate polyclonal cell lines.

### Immunofluorescence and microscopy

Cells for immunofluorescence were seeded on poly-L-lysine (Sigma-Aldrich) coated coverslips and fixed in (for the microtubules staining) with 4% formaldehyde in PHEM (60 mM PIPES, 25 mM HEPES, 10 mM EGTA, and 4 mM MgSO4) or ice-cold methanol (for the EB1 staining) for 10 min followed by plus 4% formaldehyde for in PBS 10 min.. Coverslips were washed with PBS plus 0.1% Triton X-100 and blocked in Abdil (20 mM Tris-HCl, 150 mM NaCl, 0.1% Triton X-100, 3% bovine serum albumin, 0.1% NaN_3_, pH 7.5) for 1 hour. Primary antibodies (α-Tubulin (DM1A) Mouse mAb #3873 (Cell Signaling), mouse anti-EB1 BD Transduction Laboratories™) were diluted in Abdil. The primary antibody incubation was done overnight at 4 °C. Cy3– and Cy5-conjugated secondary antibodies (Jackson ImmunoResearch Laboratories) were diluted 1:300 in PBS plus 0.1% Triton X-100. The secondary antibody incubation was performed for 1 hour. DNA was stained with DAPI. Coverslips were mounted using VECTASHIELD Antifade Mounting Medium with DAPI (Vector Laboratories). Immunofluorescence images of iKO cell lines were acquired on a DeltaVision Core deconvolution microscope (Cytiva) using a 60x/1.42NA objective, equipped with a CoolSnap HQ2 charge-coupled device camera and deconvolved where appropriate. For microtubule intensity quantification, 75 z-sections at 0.2 mm intervals were taken.

For EB3 live cell imaging, cells were seeded into 96-well glass-bottomed plates (Cellvis) and imaged using a Dragonfly 505 spinning-disk confocal microscope (Andor Technologies) equipped with a piezo Z-stage (ASI) and an iXon Ultra 888 EMCCD camera. Live cells were imaged in a humidified chamber (OKO labs) maintained at 37°C and 5% (v/v) CO_2_ using a 100× oil immersion objective NA 1.45 (Nikon, MRD01905) (pixel size 121 nm × 121 nm). tdTomato-labeled samples were imaged using a 561-nm excitation and a 594/43 emission filter. Image analysis was performed in Fiji (ImageJ, NIH) (ref).

### Protein localization

For the CARNMT1 localization we used the pCG006 plasmid (pKC254 C9orf41 cDNA (IMAGE: 4278547)). For HN1L localization we used the pCG018 plasmid (pKC54 HN1L cDNA (IMAGE: 5296086)). For HN1 localization we used a HN1-GFP stable cell line that was made using pCB48 (HN1 cDNA (IMAGE: 3140086)). For SAMHD1 localization we used pER09 plasmid (IMAGE: 81899).

### Data analysis

High-throughput sequencing reads from the pooled CRISPR screens were mapped to the sgRNA library using Bowtie. MAGeCK-RRA (Li et al., 2014) was used to generate gene scores representing the median log2 fold change in sgRNA abundance between two samples. The sgRNA-level p-value adjustment method was set to FDR, and the FDR threshold for the gene test was set to 0.05. For comparisons between cell lines, the median log2 fold changes for each endpoint sample relative to the control sample were quantile normalized, and the differential scores between drug treated and untreated were calculated using the quantile-normalized median log2 fold changes.

Quantification of fluorescence tubulin and EB1 intensity was done on unprocessed, maximally projected images using FIJI/image J. All images were acquired using the same microscope and acquisition settings for comparison. For quantification of chromosome alignment defects, cells were defined as misaligned if at least one off-axis chromosome was observed. Only cells with mature spindle structures were evaluated. The first 100 dividing cells observed were analyzed from each experimental group, for each biological replicate. For analysis of microtubule spindle intensity, a circle of 14-pixel-diameter was drawn around the spindle and the total integrated intensity was measured. Background subtraction was done by selecting a region of 3-pixel-diameter outside of the spindle and subtracting its integrated intensity from that of the spindle region. Approximately 20–25 cells were analyzed for each condition per biological replicate. For normalization of intensity levels of each iKO cell against control values, and normalization of the control value themselves, the total integrated intensity value (after background subtraction as noted above) was divided by the median control value (with background subtraction and calculated from all cells within an experiment). Statistical analyses were performed using Prism (GraphPad Software).

Single-molecule EB3 velocities were quantified from SD movies using kymographs that were generated using the ImageJ plugin KymoResliceWide v.0.4 (https://github.com/ekatrukha/KymoResliceWide). The slope of motile EB3 events in these kymographs were used to calculate EB3 velocities corrected for acquisition settings.

## Supplemental Figure Legends

**Figure S1:**
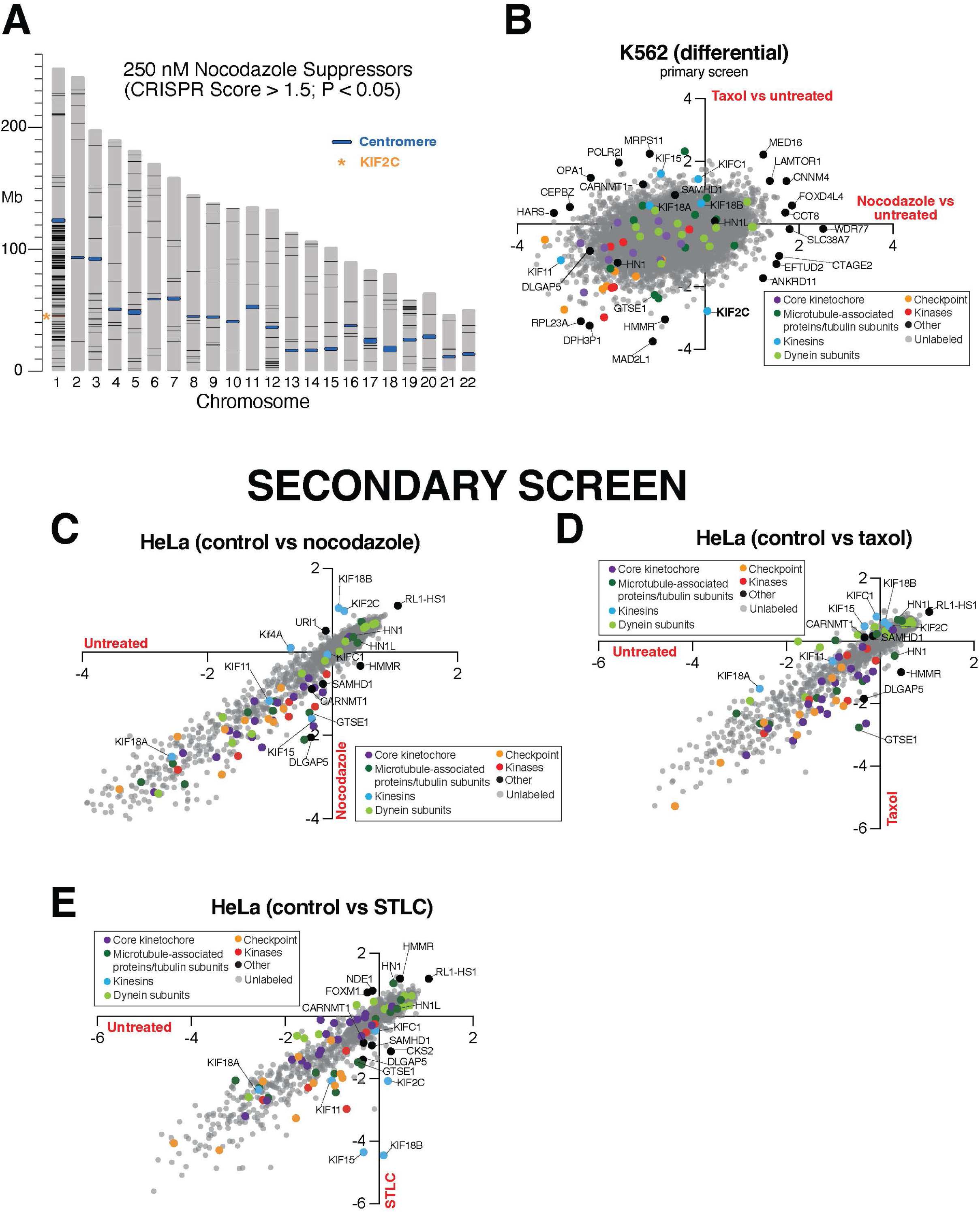
Visualization of additional screen output. (A) Autosomal start and end positions (from GRCH38.p12) of genes (black lines) with CRISPR Score > 1.5, p < .05 in 250nM nocodazole (suppressors) were obtained from BioMart and plotted in R. (B) Scatter plot illustrating the differential CRISPR scores across all gene targets in the primary screen. The differential was calculated between nocodazole and untreated and taxol and untreated K562 cell pools. (C) Scatter plot showing the CRISPR scores in untreated versus nocodazole treated HeLa cell pools from the secondary screen. (D) Scatter plot showing the CRISPR scores in untreated versus taxol treated HeLa cell pools from the secondary screen. (E) Scatter plot showing the CRISPR scores in untreated versus STLC treated HeLa cell pools from the secondary screen.

**Figure S2:**
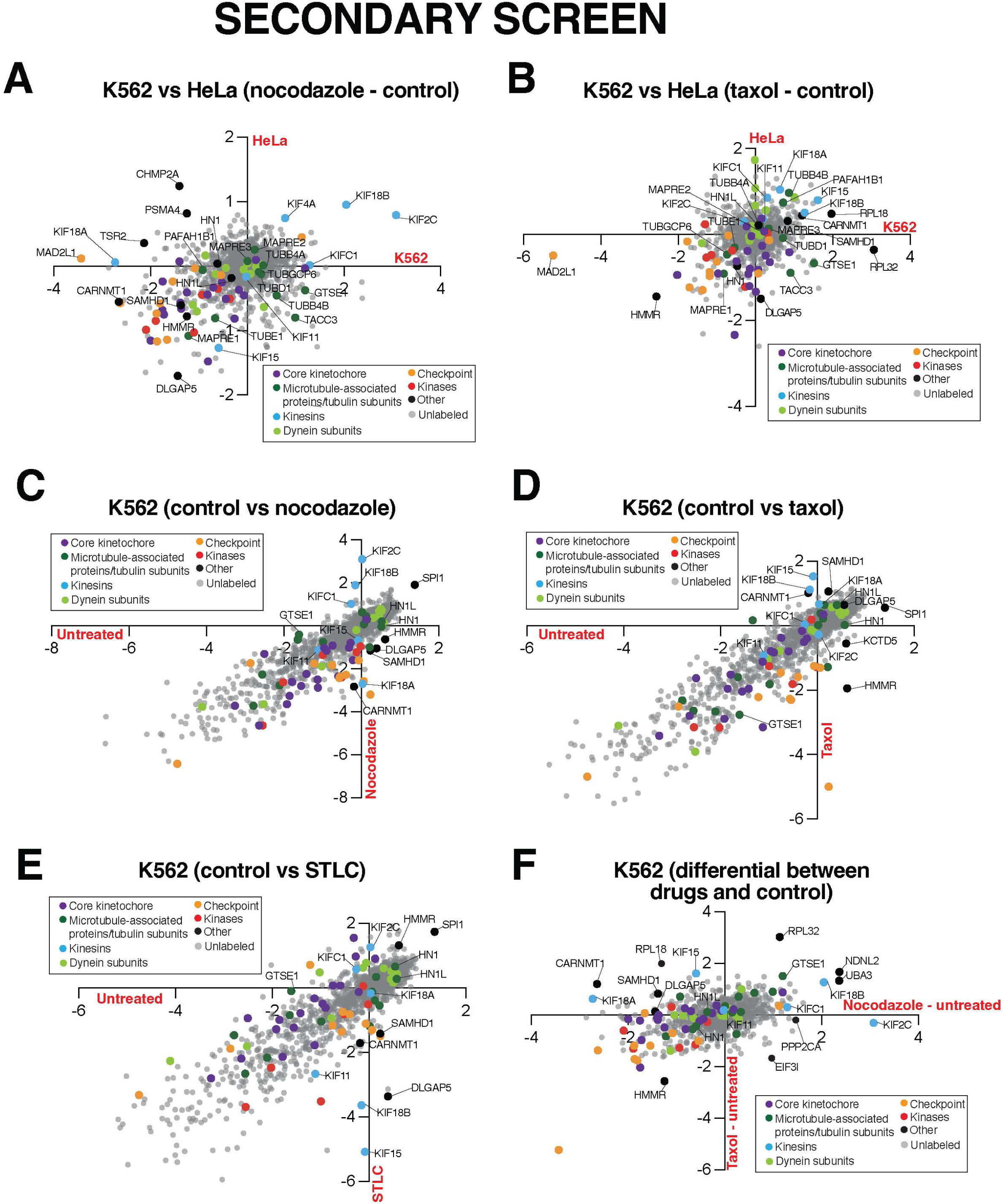
Visualization of additional secondary screen output. (A), (B) Scatter plots showing the differential CRISPR scores in treated versus untreated HeLa and K562 cells. The differential was calculated between nocodazole and untreated (A) and taxol and untreated cell pools (B). (C) Scatter plot showing the CRISPR scores in untreated versus nocodazole treated K562 cell pools. (D) Scatter plot showing the CRISPR scores in untreated versus taxol treated K562 cell pools. (E) Scatter plot showing the CRISPR scores in untreated versus STLC treated K562 cell pools. (F) Scatter plots showing the differential CRISPR scores in treated versus untreated K562 cells. The differential was calculated between nocodazole and untreated and taxol and untreated cell pools.

**Figure S3:**
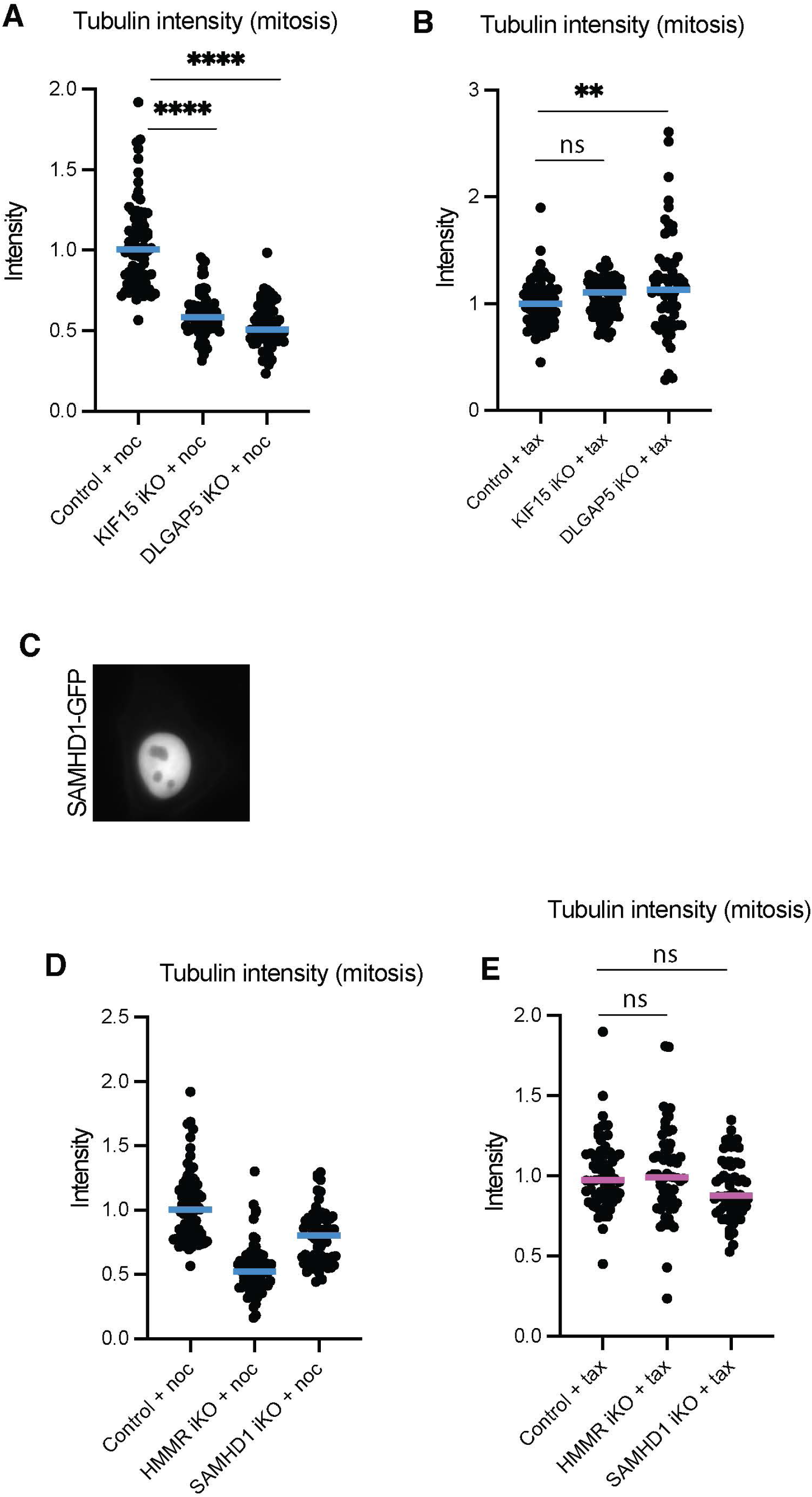
Analysis of knock-out cells of identified hits. (A) Quantification of total spindle tubulin immunofluorescence in the KIF15 and DLGAP5 iKO HeLa cells treated with nocodazole. n = 73, 70, 61 across three experimental replicates. (B) Quantification of total spindle tubulin immunofluorescence in the KIF15 and DLGAP5 iKO HeLa cells treated with taxol. n = 66, 73, 66 across three experimental replicates. (C) Representative confocal immunofluorescence images of mitotic metaphase and interphase HeLa cells showing the localization of GFP-tagged SAMHD1. (D) Quantification of total spindle tubulin immunofluorescence in the HMMR and SAMHD1 iKO HeLa cells treated with nocodazole. n = 73, 69, 67 across three experimental replicates. (E) Quantification of total spindle tubulin immunofluorescence in the HMMR and SAMHD1 iKO HeLa cells treated with taxol. n = 60, 61, 60 across three experimental replicates. Statistical tests performed: Welch’s t test (**** = < 0.0001).

**Figure S4:**
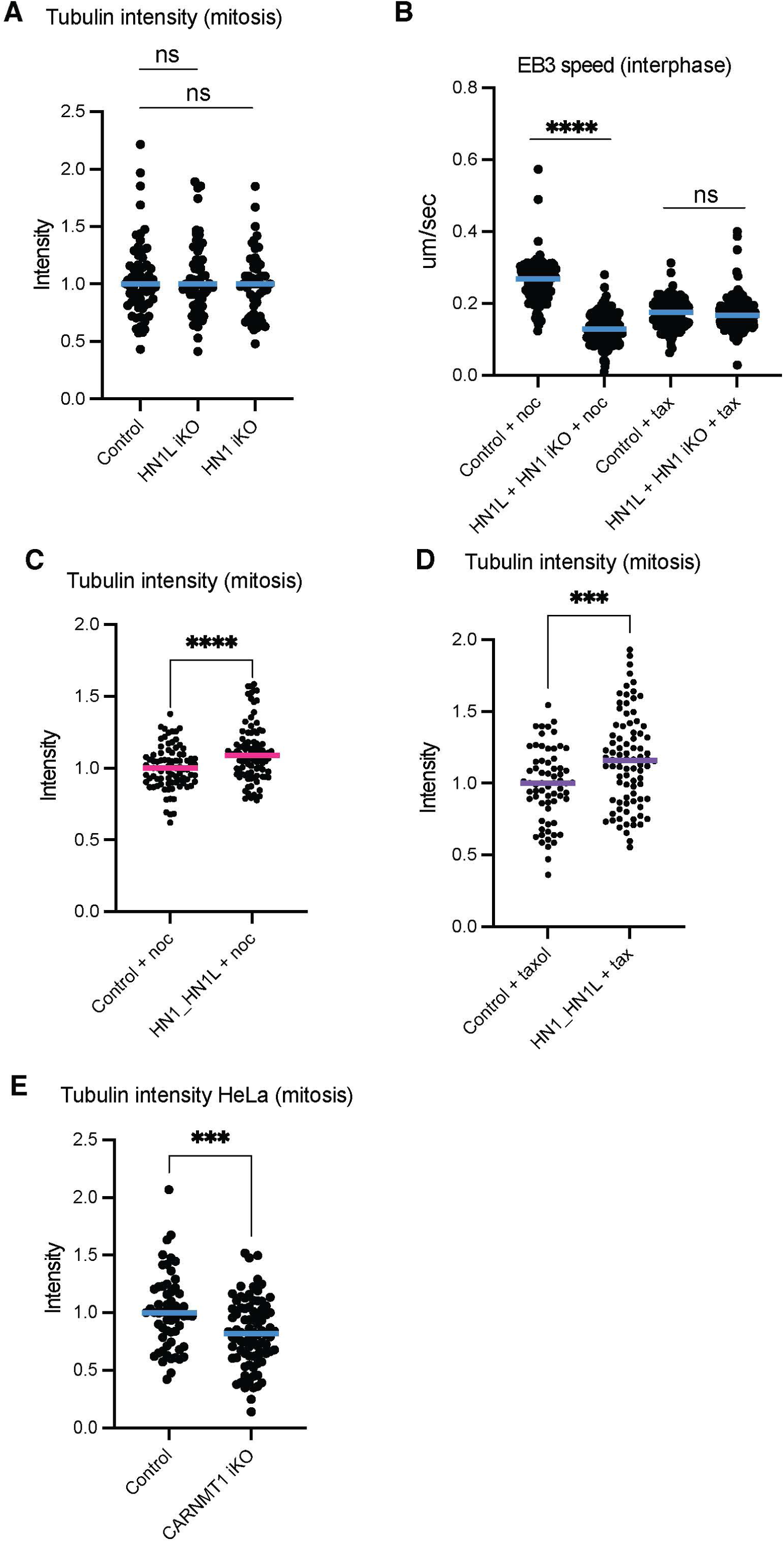
Analysis of microtubule behaviors of knock-out cells of additional hits. (A) Quantification of total spindle tubulin immunofluorescence in the HN1 and HN1L iKO HeLa cells. n = 61, 60, 52 across three experimental replicates. (B) Quantification of total spindle tubulin immunofluorescence in the HN1 and HN1L double iKO HeLa cells treated with nocodazole. n = 85, 96 across three experimental replicates. (C) Quantification of total spindle tubulin immunofluorescence in the HN1 and HN1L double iKO HeLa cells treated with taxol. n = 86, 66 across three experimental replicates. (D) EB3 speed quantification in HN1 and HN1L double iKO HeLa cells treated with nocodazole or taxol. n = 146, 153, 127, 135 kymographs, n = 28, 25, 32, 27 cells across three experimental replicates. (E) Quantification of total spindle tubulin immunofluorescence in the CARNMT1 iKO HeLa cells. n = 58, 64 across three experimental replicates. Statistical tests performed: Welch’s t test (**** = < 0.0001).

## Supplementary Tables

Table S1. Genome wide CRISPR screen sgRNA sequences, counts, and gene scores

Table S2. Secondary screen CRISPR screen sgRNA sequences, counts, and gene scores

